# “Poly (A) Binding Protein 2 is critical for stem-progenitor differentiation during regeneration in the planarian *Schmidtea mediterranea*.”

**DOI:** 10.1101/2024.04.11.588998

**Authors:** Namita Mukundan, Nivedita Hariharan, Vidyanand Sasidharan, Vairavan Lakshmanan, Dasaradhi Palakodeti, Colin Jamora

**Author notes:** Corresponding Authors Colin Jamora, Department of Life Science, Shiv Nadar Institution of Eminence, Gautam Buddha Nagar, India, Namita Mukundan, Institute for Stem Cell Science and Regenerative Medicine (inStem), Bengaluru 560065, India. Request for reprint to be addressed to Colin Jamora, Department of Life Science, Shiv Nadar Institution of Eminence, Gautam Buddha Nagar, India.

## Abstract

Post-transcriptional regulation has emerged as a key mechanism to regulate stem cell renewal and differentiation, which is essential for understanding tissue regeneration and homeostasis. Poly(A)-binding proteins are a family of RNA-binding proteins that play a vital role in post-transcriptional regulation by controlling mRNA stability and protein synthesis. The involvement of poly(A) binding proteins in a wide range of cellular functions is increasingly being investigated. In this study, we used the regenerative model organism planarian *Schmidtea mediterranea*, to demonstrate the critical role of poly(A)-binding protein 2 (PABP2) in regulating neoblast maintenance and differentiation. A deficit in PABP2 blocks the transition of neoblasts towards immediate early progenitors, leading to an enhanced pool of non-committed neoblasts and a decreased progenitor population. This is reflected in variations in the transcriptome profile, providing evidence of downregulation in multiple lineages. Thus, insufficiency of PABP2 resulted in defective formation and organization of tissue leading to abnormal regeneration. Our study reveals the essential role of PABP2 in regulating genes that mediate stem cell commitment to early progenitors during tissue regeneration.

## Introduction

Highly regenerative planarians are an exceptional model system to elucidate in vivo stem cell functions. Planarians’ regenerative capability owes to their adult pluripotent stem cell population, called clonogenic neoblast, which can differentiate into multiple cell lineages of the organism (Wagner et al 2011). Neoblasts respond to injury with a body-wide increase in proliferation rate. In response to tissue loss, a second phase of proliferation occurs locally at the wounded region critical for the formation of undifferentiated tissue known as blastema ^1^. Several regenerative clues such as signals from injury, positional information and extracellular matrix are crucial for the expression of cell type specific differentiation marks ^1–3^. In planarians, muscle fibers express positional controlling genes in a spatiotemporal manner to provide positional information helping in stem cell fate determination ^4^. It has been shown that neoblast at S, G2 and M cell cycle phases expresses fate specific transcription factors (FSTF) followed by asymmetric division to form early progenitors (specialized neoblast) and noncommitted neoblast ^5^. Some of the specialized neoblasts can undergo asymmetric division to give rise to daughter neoblasts with non-identical fate specifying factors ^5,6^. Perturbation in any of these processes including proliferation, differentiation mediated by asymmetric division or external clues can lead to defective regeneration and tissue turnover.

Studies have shown that neoblasts undergo massive changes in gene expression during the process of proliferation and differentiation. RNA binding proteins are critical for determining the functionality of transcripts thereby regulating gene expression. Several genes encoding RNA binding proteins are enriched in the neoblast population suggesting a prominent role for post-transcriptional regulation in stem cells during planarian regeneration ^7–9^. Post-transcriptional processes like alternative polyadenylation and poly A tail length modulation govern mRNA stability through intrinsic sequence alterations. Our previous study identified homolog for CPSF, CsTF, CFI, CFII, and the cis sequences binding sites in planarians, critical for polyadenylation and their regulation^10^. The Polyadenylate stretches at the 3’ UTR of mRNAs entail a high affinity to highly conserved RBPs called Poly A Binding Proteins Nuclear (PABPN) in vertebrates i.e. Poly A Binding Protein 2 (PABP2) in invertebrates. In mouse models and human cell lines, PABP2 mRNA and protein expression levels are lower in skeletal muscles at homeostasis. Interestingly, an increase in PABP2 is observed during skeletal muscle regeneration which emphasizes the need for this protein during tissue regeneration and repair ^11^. However, the role of PABP2 in modulating stem cells functioning during regenerative processes is unknown. PABP2 is a multi-functional protein involved in several molecular processes, it has been shown to interact with RNA polymerase II suggesting a possible role in transcriptional regulation ^12^. It is also identified as a molecule involved in alternative polyadenylation ^13–15^, RNA hyperadenylation ^16^, and regulation of lncRNA ^17^. The molecular mechanisms of PABP2 have been extensively studied whereas its implication in stem cell functioning is yet to be defined.

We identified Smed *pabp2* (dd_Smed_v6_21160_0_1) enriched in neoblast with a single RNA recognition domain that shows 45% sequence similarity with human PABPN. Here, we report a novel role for PABP2 in stem cell differentiation during tissue regeneration. Smed *pabp2* knockdown leads to altered phenotypes in both regenerative and homeostatic conditions. Further analysis showed that PABP2 is critical for stem cell regulation, and its downregulation restricts fate commitment during regenerative processes.

## Materials and methods

### Planarian culture

*Schmidtea mediterranea* (sexual strain) were maintained at 18-20°C in the incubator (SANYO MIR 554). The culture was maintained in 1X Montjuich salt media (1.6mM NaCl,1mM CaCl_2_,0.1mM MgCl_2_,1mM MgSO_4_,0.1mM KCl,1.2mM NaHCO_3_; pH 7) in milli Q water (Cebrià and Newmark 2005). The media was filtered using 0.45um vacuum filters (Membrane filter Millipore_654). Stock animals were fed twice per week with beef liver paste and the experimental animals were starved for 7 days prior to the experiment. The experimental animals were maintained in petri dishes (120mm) and were cleaned every alternative days.

### RNA extraction

Trizol (Invitrogen, 15596026) reagent was used for RNA extraction. The tissue for RNA extraction was collected in 1.5 ml Eppendorf tubes and incubated with 300ul of Trizol at - 80°C for 1 hour. 1/5^th^ the total volume of chloroform was added and mixed by inverting and kept on ice for 10 minutes. Subsequently the sample with the reagents were centrifuged at 21,000g for 20 minutes at 4°C. The upper most aqueous layer was collected into a fresh 1.5 ml Eppendorf tube without disturbing the interface. An equal amount of pre-chilled isopropanol was added and mixed with the previous collected solution and incubated at -20°C for 30 minutes. Following incubation, RNA was pelleted at 18,000g for 20 minutes at 4°C. The supernatant was discarded and the pellet was washed twice with 70% ethanol (prepared in nuclease free water from Invitrogen; code-10793837) by centrifuging at 18,000g for 10 minutes at 4°C. During RNA extraction from sorted cells 1ul of glycoblue (Invitrogen, AM9515) was added to visualize the RNA pellet. The washed RNA pellet was air dried and reconstituted with nuclease free water (Invitrogen_10793837) and stored at -80°C.

### cDNA preparation

1ug of extracted RNA was used to prepare cDNA using SuperScript II Reverse Transcriptase kit (Invitrogen 18064022), oligo (dT) 12-18 Primer (Invitrogen 18418012) and RNaseOUT Recombinant Ribonuclease Inhibitor (Invitrogen 10777019). The reaction was set up as per the protocol manual. The RNA in the reaction mix was digested using RNase H treatment (Invitrogen 18021014).

### Gene cloning

Gene specific primers were used to amplify specific genes (FP: AATCAATTGCATTTTTTATATCTTTAGC, RP: ATCTATCCAATTATTACTCATAACAAAAC). LA Taq DNA Polymerase (Takara RR002C) was utilized for PCR amplification of targeted genes. Purified PCR products were inserted into the vector using TA cloning kit as per their protocol manual (Invitrogen K207040). The ligated products were transformed into competent DH5α cells (Escherichia coli) and screened for blue and white colonies under a Kanamycin resistant background. Plasmids were extracted from the selected colonies and sequence confirmed through sequencing with M13 primers (Invitrogen; N52002 and N53002).

### RNA interference (RNAi)

pabp2 and gfp (used as control) dsRNA was prepared from gel eluted (Promega, A9282) PCR product using T7 RNA polymerase (NEB, M0251L) as described previously ^18^. Further the product was treated with DNase-1 (Sigma, 1014159001) and RNA was precipitated with 4M Lithium chloride and 100% ice cold ethanol. dsRNA synthesis was confirmed by running 1ul of the purified product on 1.5% agarose gel with 0.5 µg/µL ethidium bromide (Amresco, X328) in TBE (90 mM Tris/borate/2 mM EDTA; pH 8.0). The concentration of RNA was assessed by comparing the known nucleic acid concentrations of the DNA ladder bands (NEB, N3232L). Microinjection protocol was followed as mentioned previously ^19^ using Nanoject II injector (Drummond Scientific Company, Broomall, PA, USA, 3-000-204) for prepharyngeal injection of ds RNA. Three shots of 69nl (stock concentration of dsRNA-1ug/ul) were administered for three consecutive days followed by recovery time of 4 days. This regimen was followed and animals were cut into two on the 11th day and checked for phenotypic defects. In homeostasis experiment injection regimen was followed for 28-30 days.

### Whole Mount Immunostaining

Animals treated with 2% HCl and fixed using Carnoy’s fixative (60% ethanol, 30% chloroform, 10% glacial acetic acid) were stored at -20°C for at least an hour and was rehydrated before bleaching in hydrogen peroxide (20% H_2_O_2_ in methanol). The animals were incubated in blocking solution (10% horse serum in PBSTx) followed by primary antibody (anti arrestin (a kind gift from Agata lab, Japan)/ tmus (a kind gift from Sanchez lab, Stowers Institute, U.S.) / anti phospho-histone H3 ser10 (abcam 47297)/ anti-acetylated Tubulin (Sigma) incubation overnight at room temperature or 2-4 hours at room temperature at dilutions of 1:5000 (anti arrestin), 1:100 (tmus), 1:100 (anti phospho-histone H3 ser10) and 1:1000 (anti-acetylated Tubulin). Anti PIWI1 antibody was raised against NEPEGPTETDQSLS antigen in rabbit as previously described^20^. Animals were washed with PBSTx, and stained with secondary antibody (1:400 anti-mouse and rabbit conjugated with both Alexa fluor 488 and Alexa fluor 546, molecular probes) incubated overnight at 4°C or 2-4 hours at room temperature. Animals were then washed and stained with Hoechst (33342). The animals were washed, mounted on slides in mowiol (Sigma, 81381) containing dabco (Sigma D2780) and stored in dark at 4°C.

### RNA probe preparation and fluorescence in situ hybridization

Plasmid with the requisite gene sequence was linearized using not1 or hind III (New England Biolabs) restriction enzyme and was used as template. Digoxigenin-UTP RNA labelling mix (Sigma, 11277073910) or dinitrophenol-UTP RNA labelling mix (Perkin Elmer NEL555001EA), SP6/T7 (Roche; 10810274001 / Invitrogen; AM2718) polymerase along with template was used for riboprobe synthesis and RNase free DNase (NEB M0303S) was used for template digestion. RNA was purified using Bio-Rad spin mini column (7326830) as per manufacturer’s instruction. Whole-mount in situ hybridization and double fluorescence in situ hybridization was done as per the previously explained procedure ^21,22^.

### Image acquisition and quantification

Olympus SX-16 stereomicroscope was used to obtain darkfield images. To acquire confocal images FV 3000 laser scanning microscope (Olympus) with Olympus Flow View software. Image processing and quantification were done using ImageJ software (https://imagej.nih.gov/ij).

Quantification of wi1 and pabp2 colocalization (n=7) was done manually by counting double positive cells from images captured randomly across the whole animal using Fiji software. Blastemal size quantification was done by comparing blastemal area to total surface area (n=6). Muscle fibre thickness was calculated as an average diameter of three random locations per muscle fibre (total number of fibres measured from a single worm =30) (n=5). Protonephridial intensity per frame was calculated using imagej (n=6). Single fluorescent insitu hybridization was quantified manually by comparing the number of positive cells across total dapi positive cells per frame (*myoD* (n=6)*, nkx1.1* (n=6) and *agat* (n=6) respectively). H3P analysis was done by comparing the number of positive cells to the whole worm area (2dpa,n=10; 4dpa,n=5; 7dpa,n=3). For WI1/*wi1* colocalization analysis, the number of cells positive for *wi1*, WI1 and both (*wi1* and WI1) was manually counted (n=6).

### Statistical analysis

GraphPad Prism software was used for data analysis. Students t test was used to compare control and experimental animals and P<0.05 was considered significant. Experiments were conducted in biological replicates.

### Single cell transcriptome analysis

We used a single-cell transcriptome dataset published from Peter Reddien’s laboratory^23^; GEO accession number GSE111764, to extract the cells that express *pabp2*. We used the data matrix submitted in the sequence read archive (SRA) to extract only the cells that express *pabp2*. We reanalysed the single-cell data as described by Ross and colleagues (Ross et al., 2018; BioProject accession number PRJNA432445). We used Seurat (https://satijalab.org/seurat/) to analyse the single-cell transcriptome for the cells that express *pabp2* mRNA^24,25^. Based on the markers from the single-cell transcriptome dataset^23^, we classified the uniform manifold approximation and projection (UMAP) clusters as cell types. We used Log Normalize method of Seurat to normalize the dataset, which was further scaled (linear transformation) using Seurat. This scaled value was further log-transformed and plotted as a heatmap for genes of interest. We used R ggplot2 GMD and heatmap.2 to derive all the plots.

### Phylogenetic analysis

We aligned panarian smed PABP2 sequence with known PABP sequences from other species (obtained from NCBI and GenBank) using MacVector and constructed a maximum likelihood phylogenetic tree.

### Transcriptome analysis

RNA was extracted from the anterior and posterior blastema of control and *pabp2* KD animals at 0 day post amputation and 3 day post amputation. NEBNext Poly(A) mRNA Magnetic Isolation Module (Catalog no-E7490L) was used for poly A selection and NEBNext® Ultra™ II Directional RNA Library Preparation with Sample Purification Beads (Catalog no-E7765L) was used for transcriptome library preparation. Sequencing was done using NovaSeq 6000 platform using SP flowcell with 2×50bp sequencing read length. All the samples were sequenced in biological replicates. Post sequencing, 11 to 16 million paired-end (2 * 50 bp) reads were obtained. Adapters were trimmed from the reads using cutadapt v2.10 (-a AGATCGGAAGAGCACACGTCTGAACTCCAGTCA -A AGATCGGAAGAGCGTCGTGTAGGGAAAGAGTGT -u 2 -U 2). The trimmed reads were mapped to the *Schmidtea mediterranea* transcriptome ^26^ using hisat2 v2.1.0 (--rna-strandness R). The mapped reads (76-80% mapping percentage) were counted using feature Counts v2.0.0. DESeq2 v1.40.1 was used to perform the read count normalization and differential expression analysis. The plots were generated in R v4.3.0. Different planarian cell-type markers were obtained from the available single-cell transcriptome data ^23^. The sequencing data reported in this manuscript has been deposited at NCBI-Sequence Read Archive (SRA) with project ID : PRJNA1031933.

## Results

### Smed *pabp2* shows enrichment in neoblast and epidermal cell population

We investigated the expression of *pabp2* across different cell types using single cell transcriptome data (https://radiant.wi.mit.edu/app/) and identified predominant expression of *pabp2* in neoblast (35%) which was followed by the major epidermal cluster (21%), cathepsin^+^ cells (11%) and intestinal population (8%) (Fig. 1 A,B,C). The *in vivo* expression of *pabp2* in neoblast was confirmed through double fluorescent in situ hybridisation using smed *wi1* (pan neoblast marker) and smed *pabp2* (Fig 1E). Results showed 27% of the neoblast population co expressed *pabp2*, which is comparable with the single cell data revealing 35% of neoblast co expressing *pabp2* thereby confirming the single cell data. From single cell RNA sequencing on neoblast enriched for both high smed wi1 transcript and protein expression, neoblast has been classified into 12 different classes (Nb1-Nb12). Among these classes, the tspan1^+^ population (Nb2) was shown to be clononergic neoblast (cNeoblast)^27^. Expression of *pabp2* was notably high in tspan positive cNeoblast, suggesting a crucial role for PABP2 at stem cell regulation (Fig. 1D). A second-highest enrichment of *pabp2* was found in the epidermal cluster. Maintenance of epidermal integrity is imperative for wound healing and regeneration in planarians ^28^. Altogether, *pabp2* was predominantly expressed in neoblast and epidermis suggesting a critical role at stem cell functioning and regeneration.

**Figure 1:**
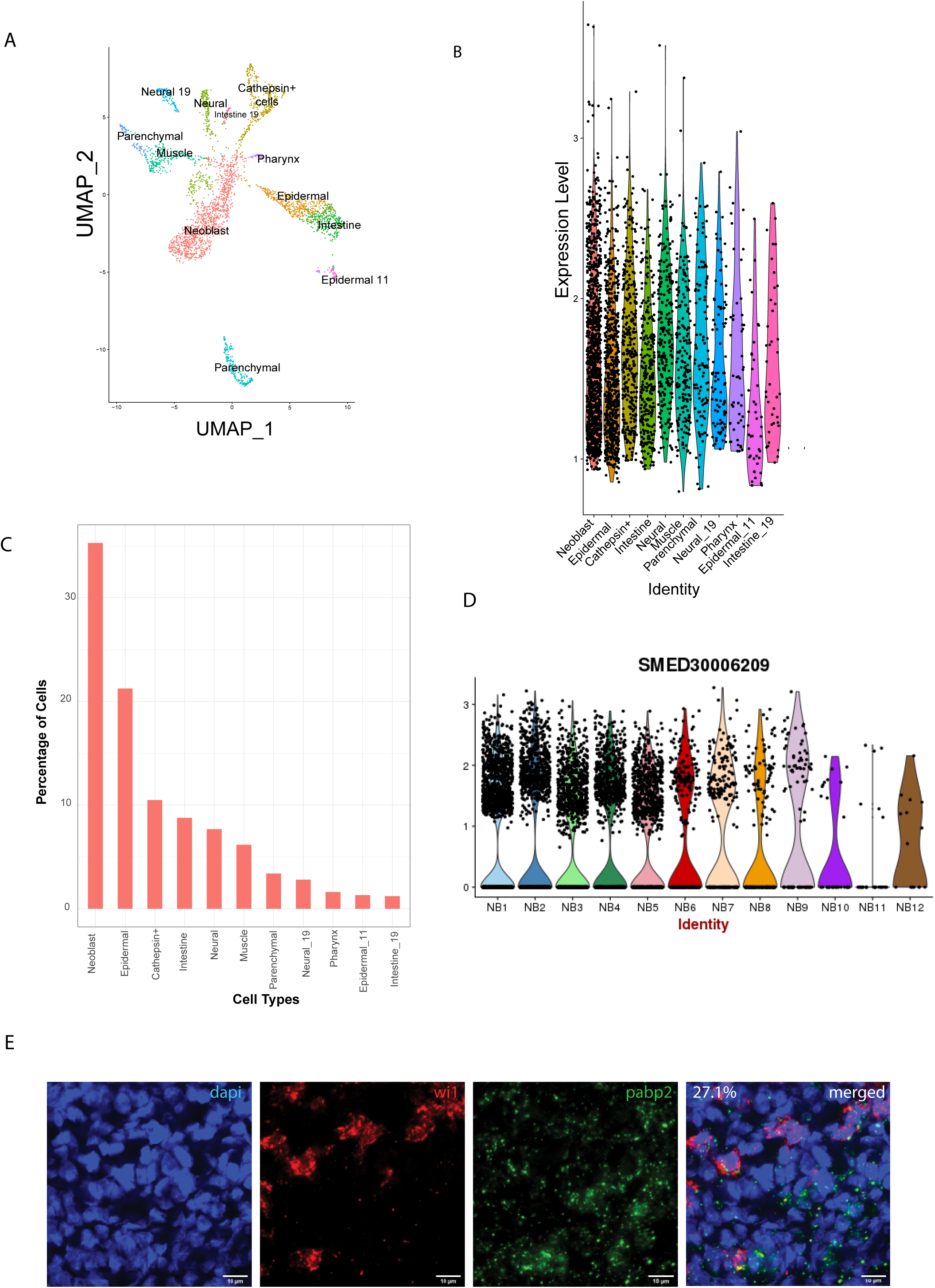
Characterization of Smed *pabp2*. A) UMAP plot obtained from single cell transcriptome data of Christopher T. Fincher *et al* 2018 showing *pabp2* distribution across different cell types. B) Violin plot depicting the expression level of *Smed pabp2* across different cell types. C) The percentage of cells expressing *pabp2* among different cell types. D) Violin plots showing the distribution of *pabp2* across 12 neoblast clusters. E) Double *insitu* hybridization showing colocalization of *pabp2* with smed *wi1*. Scale bar,10um. (n=7).

### Smed PABP2 is an essential regulator of planarian regeneration and homeostasis

Our study has pointed out an enrichment of *pabp2* among neoblasts, thereby implying an essential role for PABP2 in stem cell function. Previously, Smed-pabpc1 and Smed-pabpc2 was shown to have a role at meiotic progression and epidermal integrity respectively ^28,29^. To discern if Smed PABP2 has any functional significance during planarian regeneration and homeostasis, we knocked down *pabp2* expression. RNAi based knockdown was done through microinjection of *pabp2* and *gfp* (control) dsRNA. dsRNA was injected for three consecutive days followed by four days of recovery period. Following two rounds of injections the animals were cut in the regeneration experiment, and we observed that the knockdown animals exhibited body lesions, underdeveloped photoreceptors (Fig.2 A, B) and underwent lysis by day 20 of the injection regimen. Phenotypes were observed in 83% of animals by 8th day of regeneration. Blastema size quantification at various time points during regeneration revealed a significant (∼ 37.5%) reduction by 7th day post amputation at both anterior and posterior regenerating fragments (Fig.S2A). Under homeostatic conditions, knockdown animals showed head regression and had lesions on their body (Fig.S2B,C). 90% of animals showed defective phenotype and underwent lysis within 30 days of the injection regimen. In summary, PABP2 is necessary for tissue regeneration as well as for planarian homeostasis.

**Figure 2:**
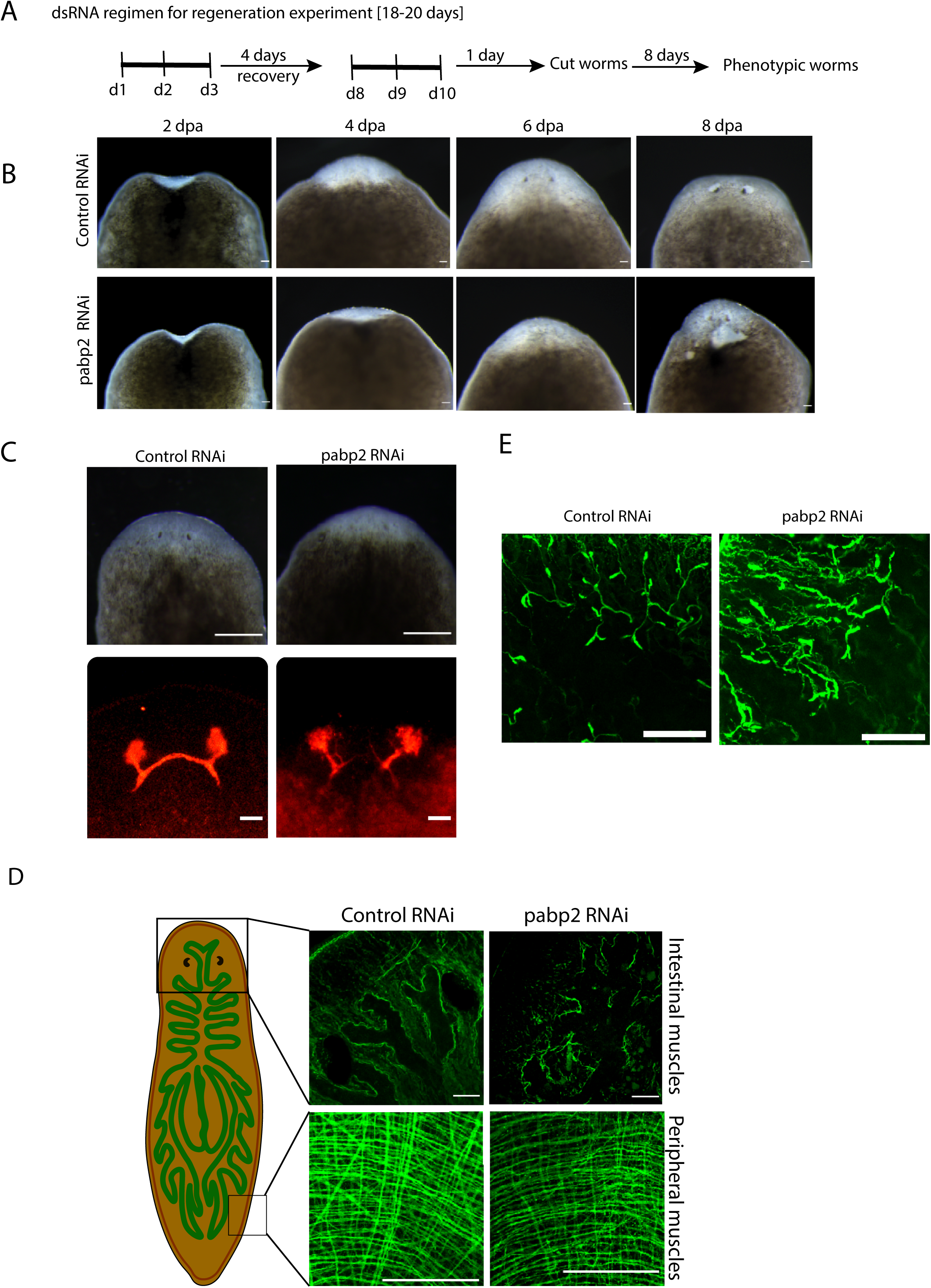
A) Schematics of dsRNA administration for regenerative RNAi experiment. B)Phenotypic animals with lesions and head regression during the course of regeneration (n=10). Scale bar,200um. C) Organization of anterior intestinal and peripheral muscles in whole mount tmus immunostaining (n=5). Scale bar,50um. D) Phenotypic animals with underdeveloped eyes. scale bar,200um and anti-arrestin immunostaining staining for photoreceptor neurons (n=5). scale bar,50um. E) Immunostaining with acetylated tubulin showing organization of protonephridia (n=6). Scale bar,50um.

### Smed*-pabp2* knockdown impairs terminally differentiated tissues

Having observed these phenotypic defects, we further focused on delineating the effect of knockdown on various tissues during regenerative processes. On the eighth day post-amputation, animals were collected and experimentally assessed for regenerative defects. The planarian photoreceptor consists of pigment cups, photoreceptor neurons (PRN) and the axonal bundle arising from the dorsally located cell body which joins together to form optic chiasma ^30–32^. Phenotypic animals were observed with reduced photoreceptor pigmentation which suggest a possible defect in eye regeneration (Fig. 2C). We investigated the organization of photoreceptors through VC1 immunostaining. We observed a lack of optic chiasma formation (Fig.2C). Previous studies reported the effect of PABP2 at muscle fiber functioning among oculopharyngeal muscular dystrophy patients ^9,33^. More specifically, in planarians, sub epidermal muscle fibers express position control genes (PCG) in regional and dynamic manner following injury thereby providing positional cues for tissue regeneration ^4^. Therefore, we examined the possibility that an impaired muscle fiber organization may be responsible for the absence of optic chiasma formation. Using tmus immunostaining, we observed that *pabp2* knockdown perturbs muscle fiber organization both at the intestinal and peripheral muscles (Fig.2D). Planarian sub epidermal myofibers are arranged in four different layers; which includes an outer circular, longitudinal, diagonal and an inner longitudinal layer ^34^. We found 33% reduction in myofiber thickness upon quantification of circular muscle fiber thickness at 8 days post amputation (Fig.S2D). Subsequently, we checked the effect of defective muscle fibers at providing positional instruction for nearby organ system. We checked the organization of protonephridia which is embedded in epithelial tissue and consist of multiple cell types that are organized to form a branched pattern ^35^ at 8 days post amputation. Anti-acetylated tubulin immunostaining revealed disorganized protonephridial branching pattern (Fig.2E). Protonephridial intensity increased by 33%, indicating a spread at protonephridial organization (Fig S2E). Taken together, we observed defects at terminally differentiated tissue which could be resulting either from an intrinsic neoblast defect failing to replace cells during regeneration or defective muscle fibers affecting the positional instructions thereby leading to disorganised organ systems.

### Knockdown of smed*-pabp2* leads to increased neoblast at the blastema

Considering the *pabp2* enrichment within the neoblast population, a depletion of PABP2 could lead to a defect of either proliferation or differentiation. One of the possibilities is a depletion of neoblast population results in decreased progenitors. A second possibility is a stable or increased intrinsically defective neoblast population affecting the later stages of differentiation. Here, we checked upon stem cell population by probing for smed *h2b* and smed *wi1* positive cells (neoblast markers) using whole mount insitu hybridization at 3 days post amputation (Fig.3A, S3A). We found a significant (p-value < 0.05) increase in Smed *h2b* and Smed *wi1* positive cells in *pabp2* KD animals suggesting an increase in the stem cell pool. In addition, we checked for mitotic neoblast (cells at G2 to M phase) using antibody against H3PS10 on 2^nd^ and 4^th^ day post amputation, which is critical for the formation of the blastema. Our analysis revealed an increased number of H3PS10^+^ cells (Fig 3 C,D) indicating an increased number of neoblast in the mitotic phase. Together, our data suggest that the neoblast maintenance might not be affected in the *pabp2* knockdown animals. However, the function of PABP2 in neoblast differentiation needs to be investigated.

**Figure 3:**
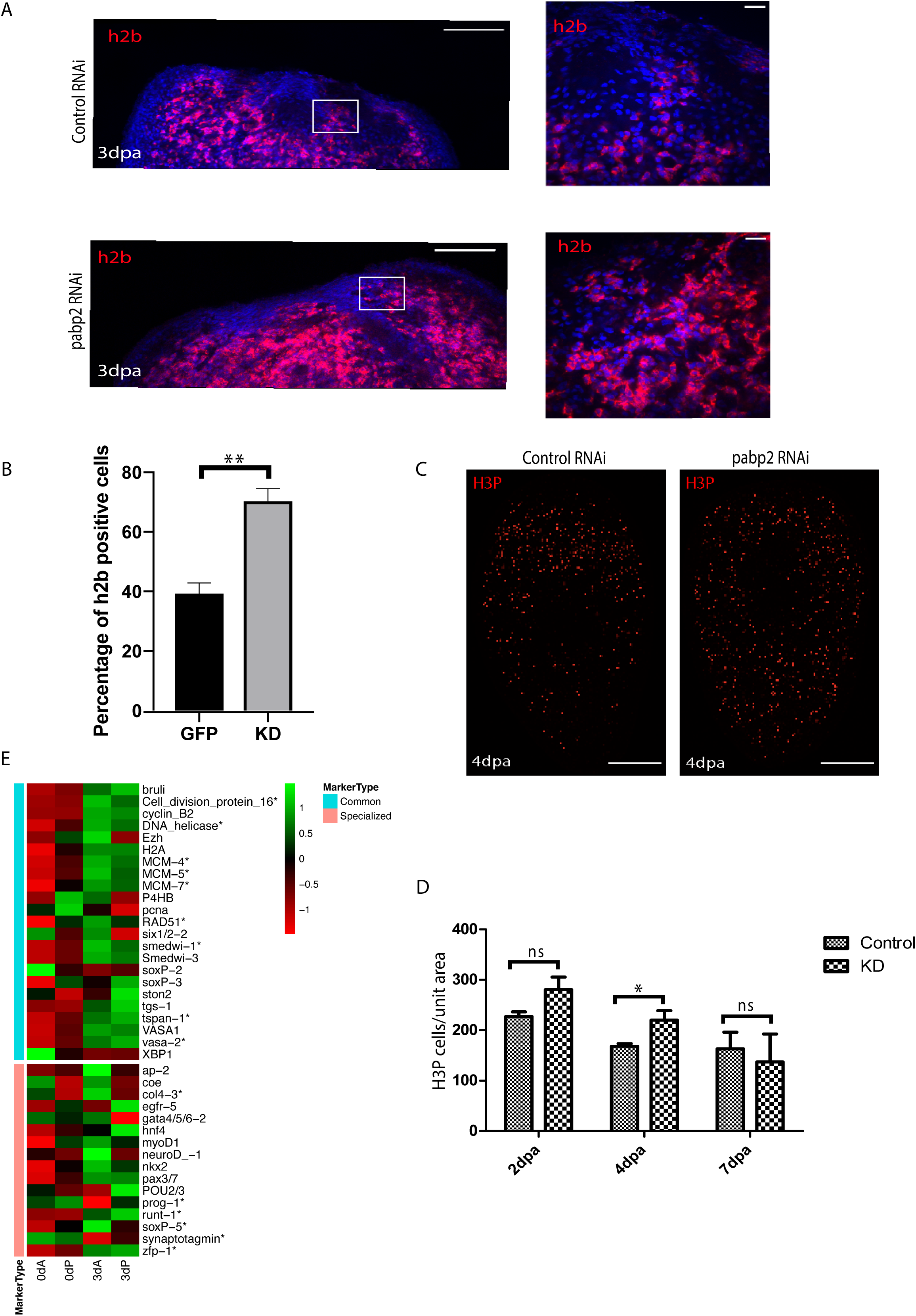
A) Confocal images of the fluorescent insitu hybridization at the regenerating surface for smed *h2b* (scale bar,100um) and the zoomed images (scale bar,20um). B) Quantification of the percentage of *h2b*^+^ cells (n=6), from the zoomed images. C) Confocal images showing H3PS10^+^ cells at 4dpa (sale bar: 50um). D) Quantification of H3PS10 at 2dpa (n=8), 4dpa (n=5) and 7dpa (n=3). E) Heat map depicting the expression of stem cell specific markers at 0 and 3^rd^ day post amputation across anterior and posterior regenerating blastema.

To further study the effect of *pabp2* KD on neoblast maintenance and its ability to differentiate, we conducted transcriptome profiling from 0 dpa and 3 dpa blastema from control and *pabp2* RNAi animals. At 3dpa, we observed a significant upregulation of neoblast specific transcripts including smed-*wi1*, *tspan*, *vasa* and cell cycle proteins like *DNA helicase*, *pcna* (Fig 3E, Table S1), which correlates with results from our in-situ hybridization experiments pointing towards an increased neoblast pool. Planarian neoblasts are a heterogeneous group of cells consisting of sigma, zeta and gamma sub populations (Wolfswinkel, Wagner, 2014). Sigma neoblasts possess broad lineage capacity and can give rise to zeta neoblasts. Zeta neoblasts characterized by the expression of *zfp-1* are essential to maintain epidermal lineage. Gamma is a subpopulation of the sigma neoblast which gives rise to the intestinal lineage. An extensive single cell transcriptome analysis from the X1 population of neoblast (dividing neoblast) identified 14 different clusters of neoblast with neoblast cluster 2 being the clonogenic neoblast. Further analysis also showed tspan-1 as a surface marker expressed on the clonogenic neoblast. Our transcriptome analysis showed increased expression of *tspan-1* in the smed-pabp2 KD animals suggesting enrichment of clonogenic neoblast in the knockdown animals. In addition, we also analyzed the transcriptome data for the expression of the early lineage markers. Most of the early lineage markers such as ap2, coe, ston2 (neuronal marker), hnf4, gata4/5/6 (intestinal marker) did not show any significant difference in expression (p-value > 0.05) or down regulated (p-value >0.05) in the knockdown animals (Fig 3E, Table S2). *zfp-1* (zeta neoblast marker) was upregulated in smed *pabp2* KD animals (Fig 3E, Table S2, S3). This further suggest that post mitotic epidermal marker might be downregulated without affecting the epidermal specific stem cell pool. In summary, our transcriptome data from the knockdown animals showed upregulation of the global stem cell specific transcripts and potential down regulation of post mitotic progeny markers indicating a potential disruption of the differentiation process.

### Smed*-*PABP2 is critical for cellular differentiation

Overall, an increase in neoblast accompanied by defects in terminally differentiated tissue could result from defective commitment of neoblast towards specific lineages. The second mitotic peak is accompanied by neoblast differentiation during lineage progression in *Schmidtea mediterranea* ^36^. In order to determine if there is a differentiation defect in the *pabp2* KD scenario a colocalization experiment probing for Smed *wi1* (*insitu* for mRNA) and Smed WI 1 (immunostaining for protein) was conducted. Smed *wi1* transcript is expressed in the neoblast whereas the corresponding protein is enriched both in the neoblast as well as the progenitors (Fig.S4A). Stem cell expresses both smed *wi1* mRNA and protein, whereas WI1 protein endures for 72 hours in post mitotic cells (Guo et al., 2006). Smed *wi1* (-)/Smed WI1 (+) cells represent the transition state from stem cell to postmitotic progeny, i.e., they mark the immediate early progeny. A change in the differentiation process can be measured by calculating the ratio of immediate early progeny with the stem cell population. An increase or decrease in this ratio will denote an increased or decreased differentiation process. Interestingly, the ratio at 3dpa was 1.33 in control animals and it decreased to 0.45 in *pabp2* KD animals, thereby indicating a significant decrease in immediate early progeny (Fig.4B,C). Taken together, these data demonstrate that a decrease in differentiation is reflected by an increase in the stem cell population.

**Figure 4:**
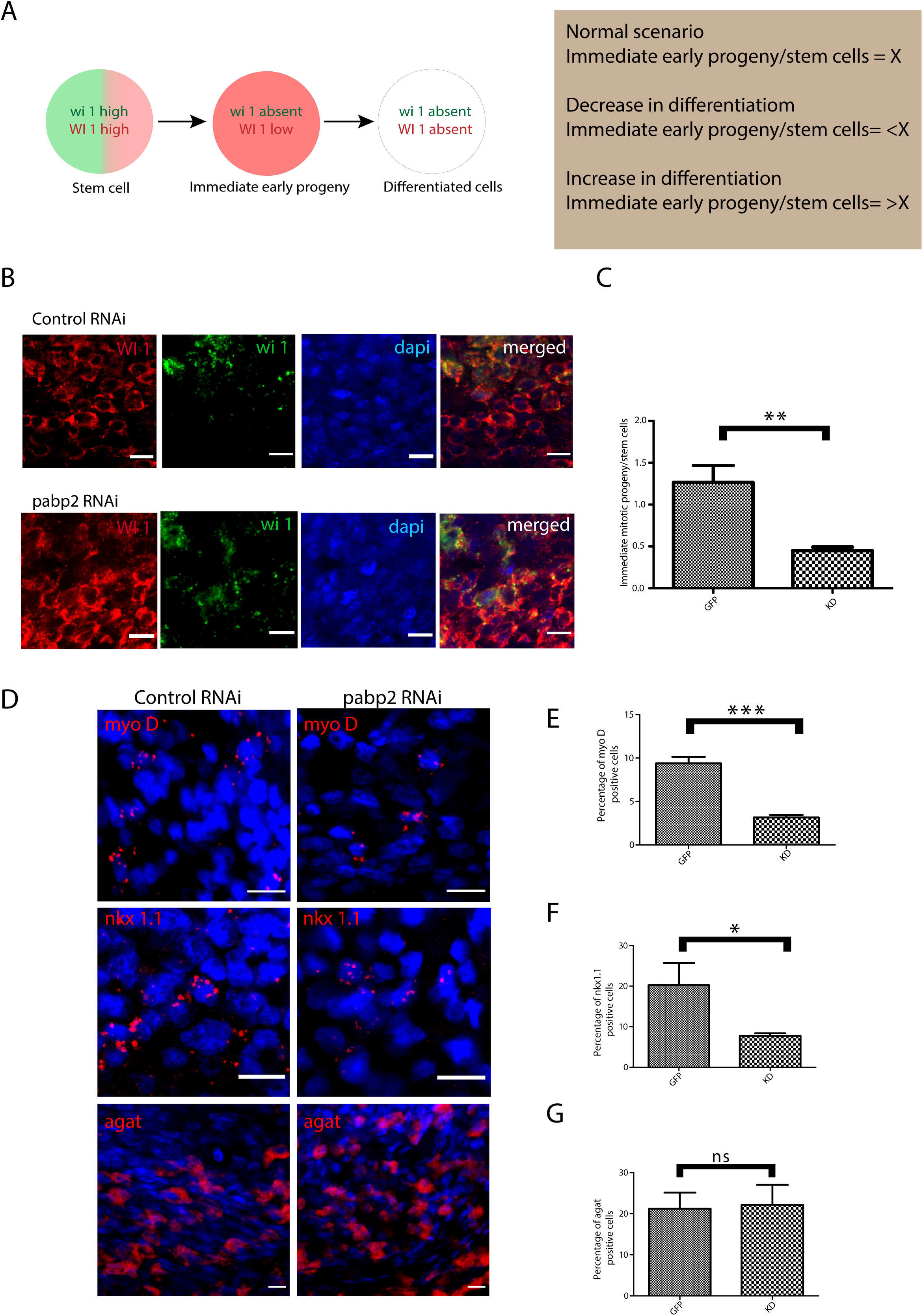
A) Schematics showing the procedure adapted for quantifying differentiation defects. B) Colocalization of smed *wi1* and smed WI1 (stem cells in green, immediate early progeny in red). Scale bar,10um. C) Graph plotted from quantified ratio of immediate early progeny to stem cell population (n=6). D) *insitu* hybridization probing for various progenitor populations and its quantifications (*myoD* - longitudinal muscle fibre, *nkx1.1* - circular muscle fibre, *agat* – epidermal cells) across control and *pabp2* KD. Scale bar,10um. E,F,G) Representation of the percentage of progenitor expressing cells compared across control and knockdown for *myoD* (n=5)*, nkx1.1* (n=6) and *agat* (n=6) respectively.

It has been shown that neoblast expresses fate specifying transcription factors required for their transition towards respective progenitors ^38^. To decipher how the progenitor populations are affected, we performed FISH by probing for markers such as myoD and nkx1.1 which mark the longitudinal and circular muscle fibres respectively and found a 60% decrease in *myoD* and 50% decrease in *nkx1.1* expressed cells (Fig. 4E,F), corroborating with the observed muscle fibre defect. Lineage specification during the epidermal differentiation process is very well studied in planarians. It is also known that zeta neoblast lineage gives rise to epidermal tissue and *agat* marks the latter epidermal progenitors ^39^. Since we found an enrichment of *pabp2* in the epidermis (Fig. 1), we quantified the epidermal progenitor population. However, there is no change in the *agat*^+^ epidermal progeny in *pabp2* KD animals (Fig.4G). This also corroborates with our previous finding that the *pabp2* KD animals showed increased expression of zfp1, a zeta neoblast marker. The lesions observed could be due to the faulty body wall muscle organization, which was evident from muscle staining in the KD animals (Fig.2D). Altogether, we found a decrease of *myoD* and *nkx 1.1* expressing cells whereas *agat*^+^ cells remain unchanged suggesting a possibility of muscle lineages being downregulated. To conclusively understand the affect of *pabp2* KD on other lineages, it is essential to study the expression of wide variety of lineage specific markers in KD animals.

### Smed-PABP2 regulates transcripts essential for major epidermal population and intestinal lineages

Transcriptome analysis was done to understand the effect of *pabp2* KD on commitment towards various cell lineages. It has been reported that most lineage specification marks are expressed by day 3 post amputation^1^. Transcriptome sequencing was conducted at 0 and 3 days post amputation blastema in control and KD animals. We investigated the expression of different cell-type markers^23^ in our transcriptome data. Among the various cell types, transcripts representing major epidermal clusters were drastically downregulated such as *egr5*, *zpuf-6*, and *vim-3* by 3dpa (Fig 5A, Table S3), which are critical for later stage epidermal differentiation (Tu et al., 2015). At 3dpa, there was a minor change in *prog* expression (log2FC -0.83 and -0.38 respectively in the anterior and posterior regenerating blastema). In the anterior and posterior regenerating blastema, we did not observe a reduction in agat expression at 3dpa (log2FC -0.05 and -0.01), and our insitu hybridization showed no change in agat^+^ cell population (Fig.4D). Together, our results suggest that epidermal lineage defects in *pabp2* KD are caused by defects in later stages of differentiation among epidermal cell types rather than early progenitors.

**Figure 5:**
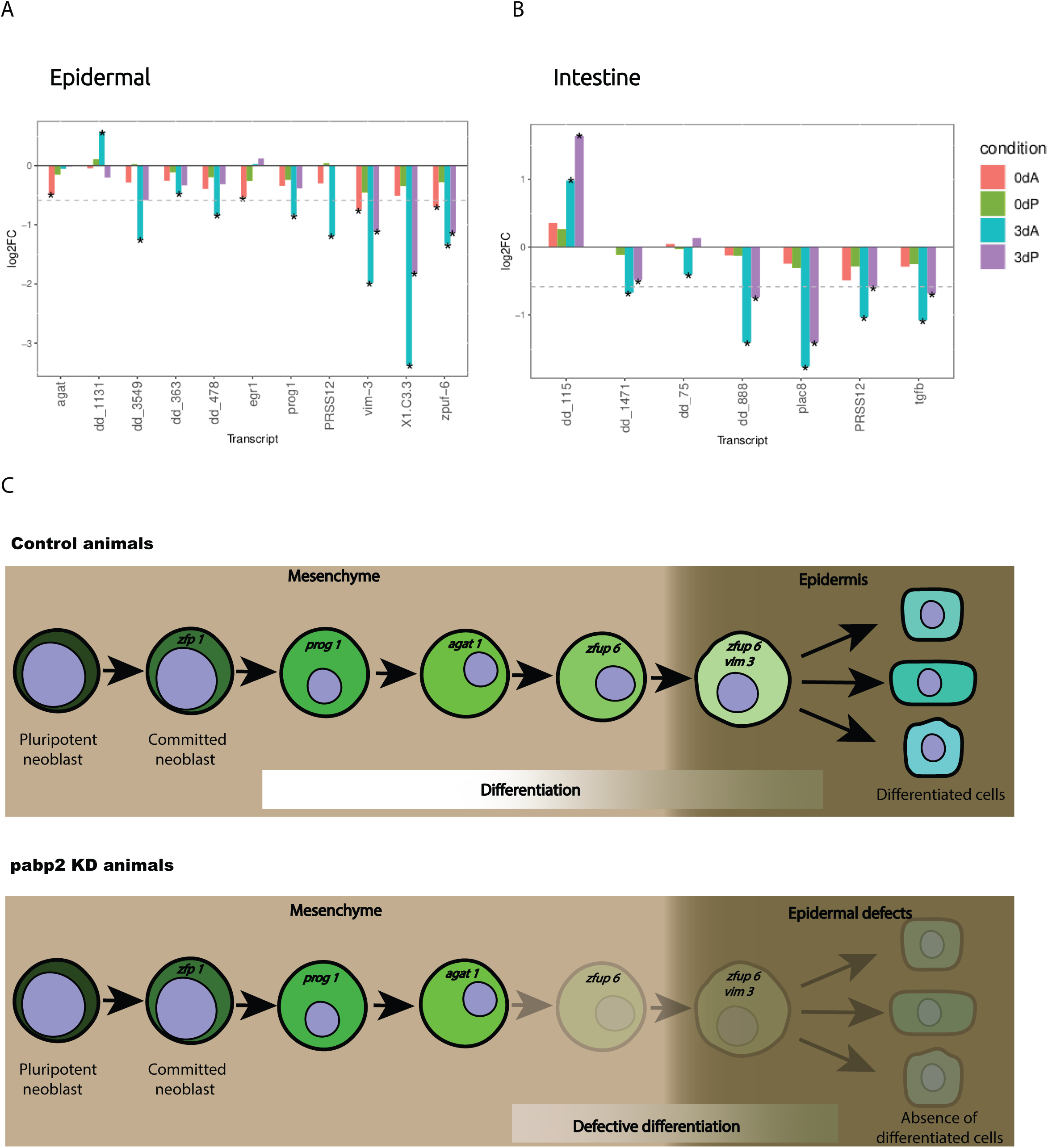
Bar plot showing gene expression variation among specific lineages. Transcripts with variations among (A) epidermal and (B) intestinal markers when compared between *pabp2* KD and control animals at various time points (as denoted by the colour bars within the plots) among anterior and posterior regenerating blastema. The asterisk indicates statistical significance (p-value <0.5) and the dotted line represents log2FC -0.58. C) Working model depicting the critical role of Smed PABP2 in epidermal differentiation.

Gamma neoblasts is a sub population of sigma neoblasts, they have the potential to differentiate towards intestinal lineage^40^. Our analysis revealed a drastic decrease in gamma neoblasts and intestinal specific transcripts across anterior and posterior developing blastema (Fig 3E,5B). Cathepsin^+^ cluster consist of heterogeneous group of cells which includes pigment cells, glial cells and several other unknown cell types ^41^. Cela1 and jag1 specific for cathepsin^+^ cells showed downregulation at 3dpa (Fig S5A, Table S3). Suggesting that among the mixed population of cathepsin^+^ cells few subtypes were affected upon *pabp2* RNAi. In the posterior developing blastema, we also observed few transcripts being downregulated among the pharyngeal cluster (Fig S5B, Table S3). Further, we also observed few muscles specific transcripts being downregulated which could possibly result from an indirect effect of suboptimal level of *papb2* (Fig S5C, Table S3). The cell types which did not show significant variation includes the parapharyngeal, neuronal, ciliated neuronal and non-ciliated neuronal lineages upon *pabp2* KD (Fig S5 D,E,F,G, Table S3). In summary, our bulk transcriptome data from the control and the KD animals overlayed on the existing single cell data showed defects predominantly in the major epidermal population, a sub set of cathepsin^+^ cell types and intestinal lineage.

## Discussion

We identified Smed *pabp2* (dd_Smed_v6_21160_0_1) enriched in neoblast. In our phylogenetic analysis, the identified PABP2 clusters with PABPN homologs from other model organisms (Fig. S1B). Our study reveals a role for PABP2 in the transition of stem cells to differentiated cell types. PABP2 insufficiency leads to enhanced self-renewal and dysregulated differentiation. Hence, a defect in PABP2 functioning affects the process of differentiation thereby leading to an insufficient progenitor population resulting in improper tissue regeneration and organization.

Cellular differentiation is the process by which committed cells undergo drastic gene regulations to become specialized cells. In human, defective differentiation can lead to several disease conditions affecting specific cell lineages such as muscle ^42^, B-cells ^43^. Here, in *pabp2* KD, an increased neoblast population along with a decrease in differentiation corroborates to a defect at lineage progression towards single or multiple cell types. We tackled the effect of PABP2 deficiency at regulating differentiation using epidermal lineages as a proxy. Planarian epidermal lineage commitment has been extensively studied, and many markers of epidermal lineage progression have been identified ^39^. The upregulation of zfp1 (committed neoblast marker) indicates the commitment of pluripotent stem cells to the epidermal lineage. Simultaneously, this is also evident from maintenance of agat^+^ cells, an epidermal lineage marker. Together, these provides evidence that the stem cell self-renewal and commitment towards epidermal lineages are unaffected. However, our transcriptome data revealed that the consequence of *pabp2* KD occurs at the latter stages of differentiation which is evident from the drastic decrease in *zpuf-6* and *vim-3* expression (late epidermal progeny markers)(Fig. 5C). This late stage differentiation defect could either be an effect of an intrinsically defective early progenitors or could result from the depletion of the late progenitor specific transcript which restricted the epidermal lineage progression. These observations point to a critical role for PABP2 in regulating cellular differentiation.

In addition to the significant reduction in transcripts specific to major epidermal population, a subset of cathepsin^+^ cell type, and intestinal populations is observed, we also observed significant decrease in the nkx1.1^+^ and myoD^+^ cells in the KD animals. This suggest that PABP2 is crucial for the regeneration and maintenance of the muscle wall. Here, it is likely that lesions observed in the KD animals could be due to the defective turnover of epidermis and muscle. Embedded in the epidermis, and surrounded by the muscle are the excretory units or protonephridia that regulate the osmotic pressure. We do not see a depletion of protonephridial transcripts, although an abrupted protonephridial branching pattern was observed. Here, the defect at protonephridia seems like a defect caused either by an epidermal irregularity or due to an indirect effect of defective muscle cells. Gamma neoblasts are a subpopulation of sigma neoblasts, which is the major stem cell population. Several gamma neoblast markers were downregulated among *pabp2* KD animals. This could be caused by sigma neoblasts’ inability to switch to the intestinal lineage, which is evident from our transcriptome data. Together, our study show PABP2 is a critical regulator for stem cell differentiation towards multiple cell lineages. Further, a detailed analysis of the impact of *pabp2* KD on other cell types is required, prioritizing intestinal lineages and cathepsin^+^ subtypes expressing *cela 1* and *jag 1*.

The current study provides insight into the role of PABP2 in regulating stem cell differentiation in planaria. However, it is necessary to delineate the molecular mechanism through which multifunctional PABP2 elicits its effects during regeneration. Developing PABP2 antibodies specific to *Schmidtea meditteranea* would be necessary to understand the direct interacting partners. This would enable us to delineate the mechanistic role of PABP2 in stem cell functioning. Studies have shown that mitochondrial dynamics regulate stem cell self-renewal and differentiation^44–46^. We observed downregulation of mitochondrial potential among *pabp2* KD animals (data not shown); suggesting a possibility of PABP2 modulating mitochondrial transcripts specific for stem cell fate determination. It is possible that PABP2 could play an inevitable role in regulating mitochondrial transcripts necessary for stem cell differentiation.

Conventionally, PABP2 is believed to be ubiquitously involved in actively transcribing transcripts. Later, a number of studies have analysed its multifaceted functions, including transcriptional regulation^12^, alternative polyadenylation^13–15^ and hyperadenylation and decay of RNA^16^. Recent studies conducted among xenopus, mouse and MEL (mouse erythroleukemia) cells have reported the role of poly A binding proteins in regulation of maternal transcript processing and erythroid differentiation, thereby, extrapolating its function to stem cell regulation^47–49^. Till date, few molecules requisite for neoblast differentiation have been identified among *Schmidtea meditteranea*, which includes Smed mex3, Smed CHD4, Smed apob^50–52^. Our study demonstrates the requirement for PABP2 during stem cell differentiation in planaria.

**Figure S1. Related to figure 1:** A) Phylogenetic tree showing the relatedness of poly A binding proteins.

**Figure S2. Related to figure 2:** A) Blastema size quantification from 1^st^ to 7^th^ days post amputation (n=6). B) Timeline showing injection regime where ds RNA was injected during homeostasis experiment. C) Homeostasis knockdown animals with lesions at the anterior region (n=7), scale bar,200um. Images were taken on the 28^th^ day from the beginning of dsRNA regimen. D) Quantification of peripheral muscle fibre thickness (three frames from a single worm where 10 fibres were measured from each of the frames. Total fibres measured from a single worm=30; n=5). E) Quantification of intensity of anti acetylated tubulin staining for protonephridia; n=6.

**Figure S3. Related to figure 3:** A) Confocal images of the fluorescent insitu hybridization at the regenerating surface for smed *wi1* (scale bar,100um) and the zoomed images (scale bar,20um). B) Quantification of the percentage of smed *wi1*^+^ cells (n=5).

**Figure S4. Related to figure 4:** A) Confocal images of the entire animal showing distribution of stem cell population (smed *wi1*-green) differentiating cell population (smed WI1-red) and immediate early progenitors (merged-yellow) (n=6); scale bar,100um.

**Figure S5. Related to figure 5:** Bar plot depicting gene expression variations. Transcripts with variations among A) Cathepsin^+^ cells B) Pharynx C) Muscle D) Parapharyngeal E) Neural F) Ciliated neurons G) Non ciliated neurons when compared between pabp2 KD and control animals during time points (as denoted by the colour bars within the plots) among anterior and posterior regenerating blastema. The asterisk indicates statistical significance (p-value <0.5) and the dotted line represents log2FC -0.58.

**Table S1. Related to figure 3:** Expression values of common neoblast markers from RNA sequencing analysis at 0dpa and 3dpa anterior and posterior regenerating blastema.

**Table S2. Related to figure 3:** Expression values of specialized neoblast markers from RNA sequencing analysis.

**Table S3. Related to figure 5:** Expression values of differentiation markers from RNA sequencing analysis.

## Acknowledgement

We are grateful for the immense technical support from the Central Imaging and Flow cytometry Facility (CIFF) and Sequencing facility (special thanks to Dr. Awadhesh Pandit) at the BLiSc Bio-Cluster. We thank Sanchez’s laboratory and Agata’s laboratory for their generous gift of tmus and VC1 antibodies. We also acknowledge the support from thermo scientific for developing WI1 antibody.

## Authors Contributions

Namita Mukundan and Dasaradhi Palakodeti conceived and designed the experiments. Namita Mukundan and Vidyanand Sasidharan performed the experimental procedures. Namita Mukundan performed the experimental analysis. Nivedita Hariharan and Vairavan Lakshmanan performed the transcriptome analysis. Namita Mukundan prepared the figures. Namita Mukundan, Colin Jamora and Dasaradhi Palakodeti drafted the article.

## List of Abbreviations

cNeoblast: clononergic neoblast
dpa: days post amputation
FSTF: fate specific transcription factors
FISH: Fluorescent insitu hybridization
MEL: mouse erythroleukemia
PABPN: Poly A Binding Proteins Nuclear
PABP2: Poly(A)-Binding Protein 2
PCG: position control genes
PRN: photoreceptor neurons

