## Supplementary figures and images for "“Poly (A) Binding Protein 2 is critical for stem-progenitor differentiation during regeneration in the planarian *Schmidtea mediterranea*.”"

### Supplemental Figure 1

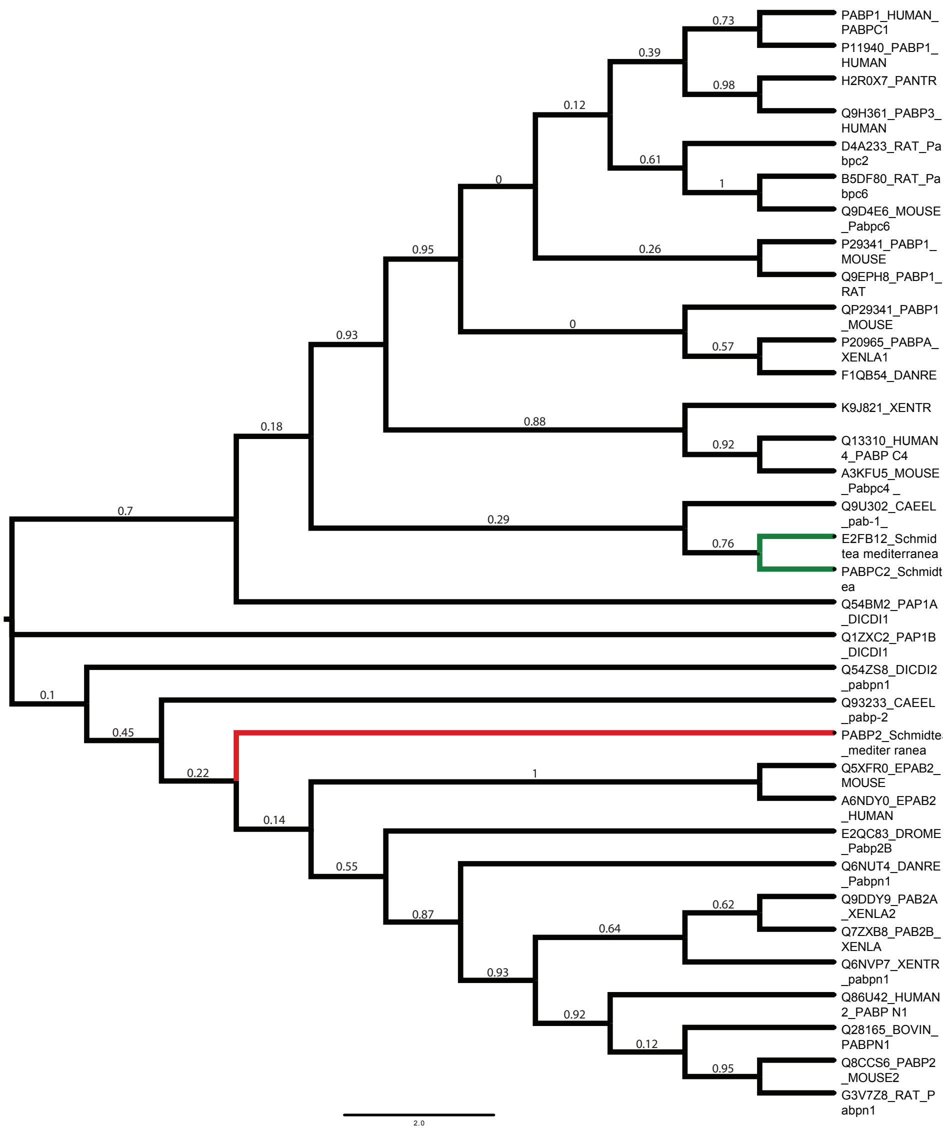

### Supplemental Figure 2

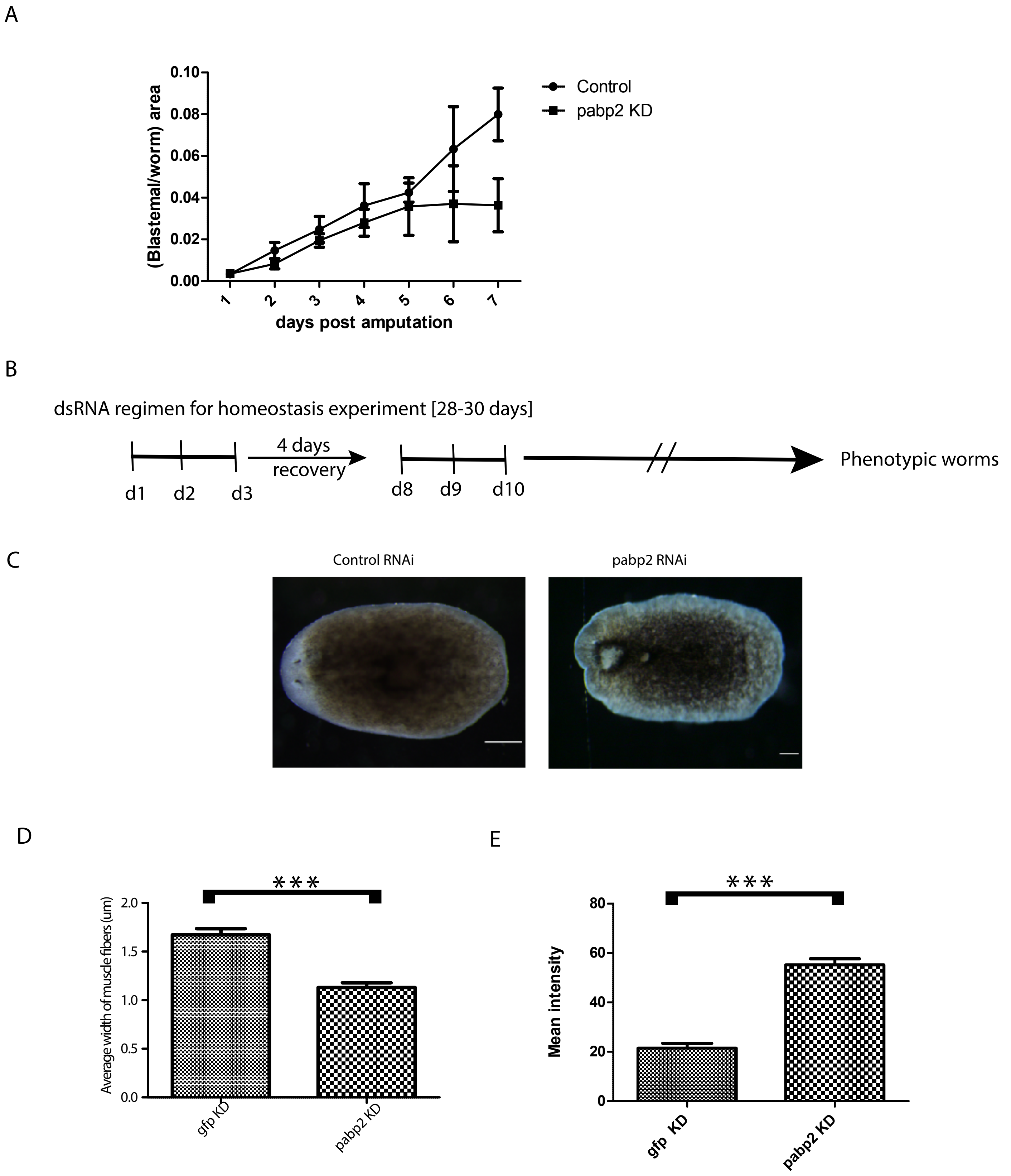

### Supplemental Figure 3

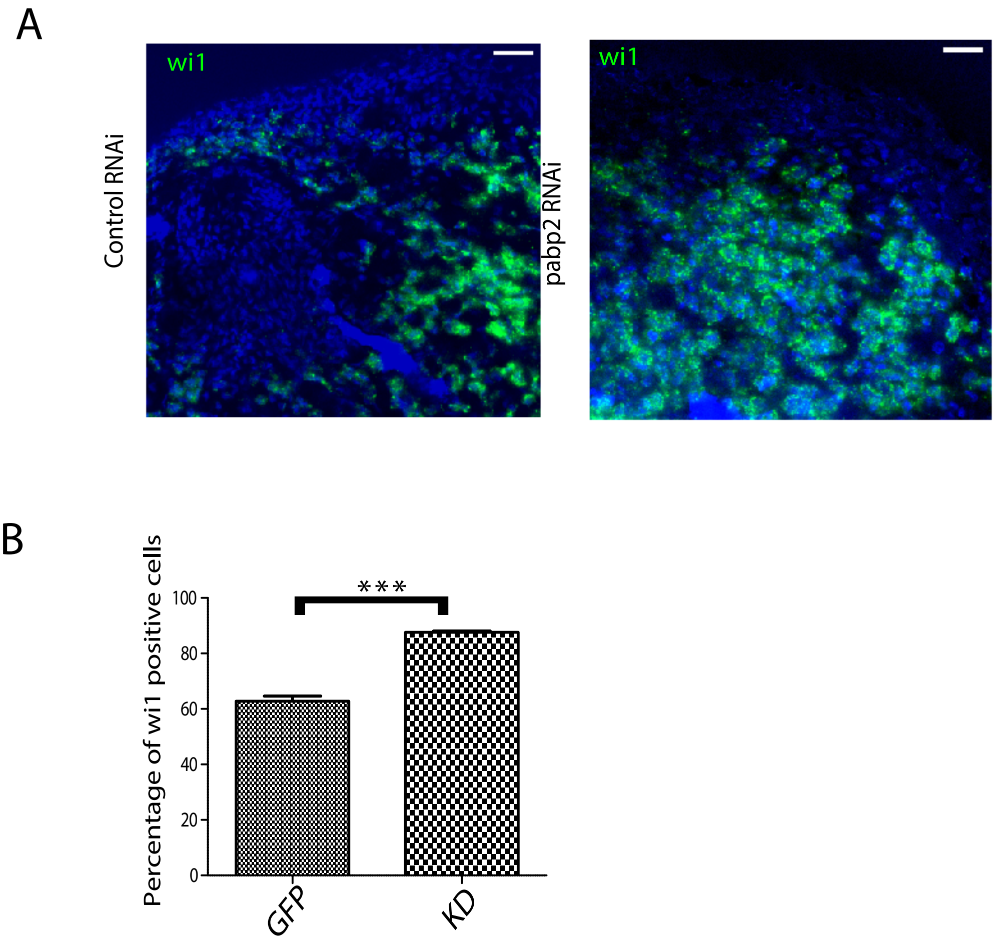

### Supplemental Figure 4

A

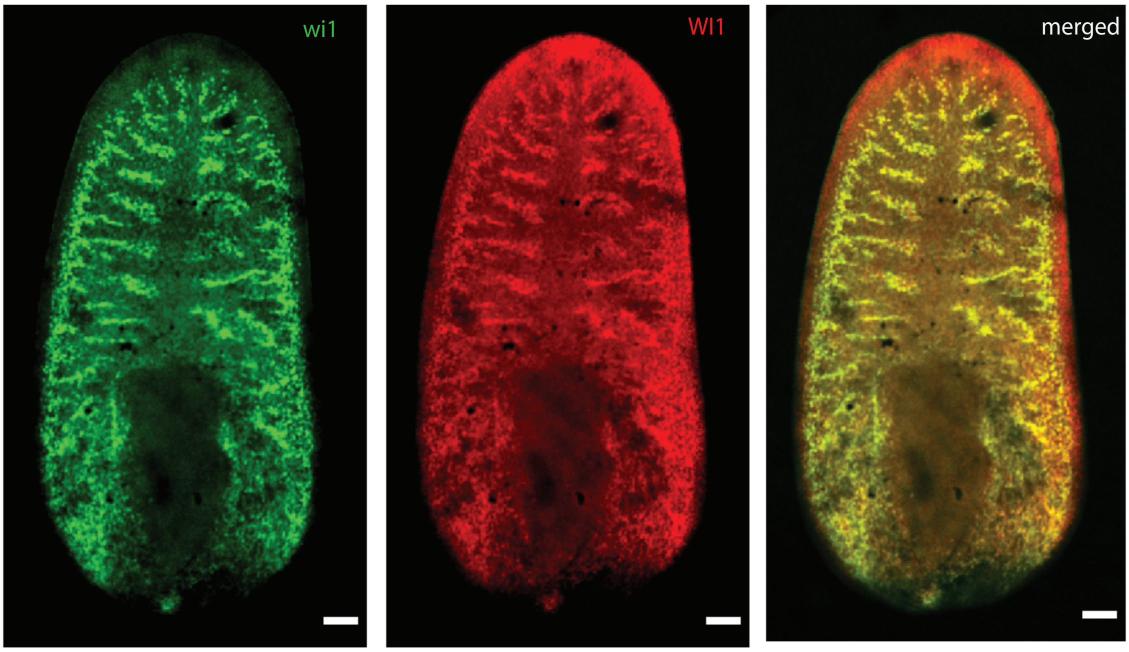

### Supplemental Figure 5

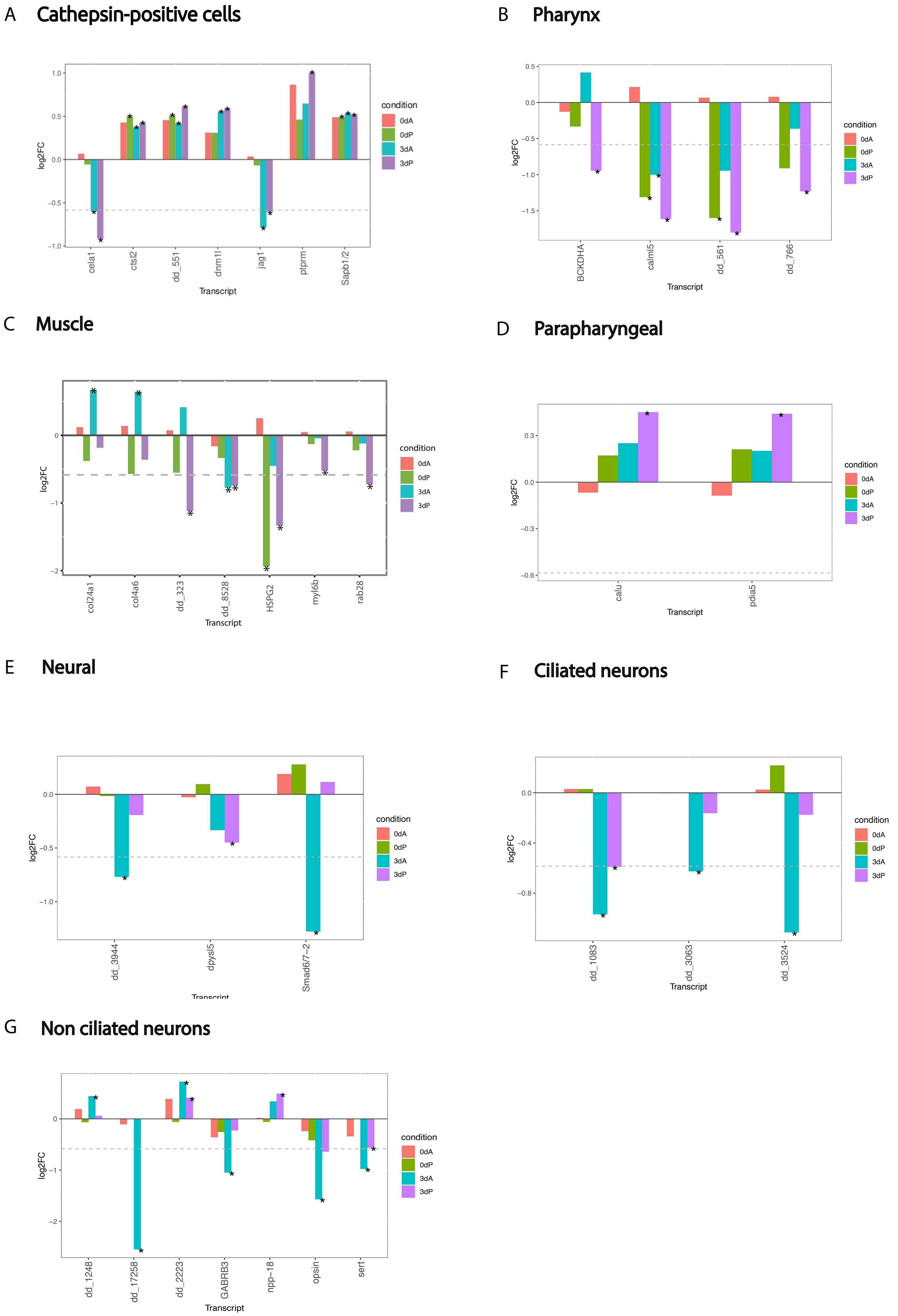
